# A sex chromosome inversion is associated with copy number variation of mitochondrial DNA in zebra finch sperm

**DOI:** 10.1101/776054

**Authors:** Ulrich Knief, Wolfgang Forstmeier, Bart Kempenaers, Jochen B. W. Wolf

**Author notes:** Address for correspondence: Ulrich Knief, Division of Evolutionary Biology, Faculty of Biology, Ludwig Maximilian University of Munich, Grosshaderner Str. 2, 82152 Planegg-Martinsried, Germany.

## Abstract

Propulsion of sperm cells via movement of the flagellum is of vital importance for successful fertilization. Presumably, the energy for this movement comes from the mitochondria in the sperm midpiece. Larger midpieces may contain more mitochondria, which should enhance the energetic capacity and hence promote mobility. Due to an inversion polymorphism on their sex chromosome *TguZ*, zebra finches (*Taeniopygia guttata castanotis*) exhibit large within-species variation in sperm midpiece length, and those sperm with the longest midpieces swim the fastest. Here, we test through quantitative real-time PCR in zebra finch ejaculates whether the inversion genotype has an effect on the copy number of mitochondrial DNA. Taking the inversion genotype as a proxy for midpiece length, we find that zebra finches with longer midpieces indeed have more copies of the mitochondrial DNA in their ejaculates than those with shorter midpieces, with potential downstream effects on the rate of ATP production and sperm swimming speed. This study sheds light on the proximate cause of a fitness-relevant genetic polymorphism, suggesting the involvement of central components of gamete energy metabolism.

**Data availability:** Supplementary data file

## 1. Introduction

Sperm morphology can have direct consequences for sperm motility and male reproductive success [1-3]. Thus, sperm competition and cryptic female choice should exert strong directional selection towards an optimal sperm design, presumably enhancing motility or energetic capacity [4]. Yet, the proximate connection between sperm design and motility remains poorly understood.

A typical sperm cell moves through the activity of its flagellum [5], which consists of the midpiece and the tail [6]. It derives the energy for its movement either through anaerobic glycolysis in the tail or through oxidative phosphorylation (OXPHOS) in the mitochondria of the midpiece [5]. Whereas mammalian species vary considerably in their use of these two energetic pathways [5], OXPHOS alone supports normal sperm motility in chicken (*Gallus gallus*; [7]), suggesting respiration in mitochondria as the main energy source [4]. In birds, the total number of mitochondria per sperm cell varies from less than 20 to more than 350 across species [6]. In oscine songbirds, the largest phylogenetic group of birds, all mitochondria fuse into a helical strand that winds along the midpiece (gyres) during spermatogenesis [8-10]. The number of mitochondria contributing to this mitochondrial syncytium is basically unknown. Because sperm midpiece length correlates with adenosine triphosphate (ATP) content in whole ejaculates across songbirds, it has been hypothesised that a larger number of mitochondria contributes to the syncytium in longer midpieces [11].

Zebra finches are polymorphic for an inversion on their sex chromosome *TguZ* that spans roughly 63 megabases and 619 genes [12, 13]. Kim *et al.* [14] and Knief *et al.* [2] have previously shown that this inversion has profound effects on sperm morphology, sperm velocity and fertilization success. Males that are heterozygous for the inversion have sperm with a long midpiece and a relatively short tail (**Figure 1a**). The total flagellum length is intermediate between those of the two homozygous groups. This design seems optimal, because these sperm swim fastest and fertilize the most eggs in a competitive environment [2]. The allele frequency of the ancestral inversion haplotype in a wild Australian population (59.6%; [12]) might be explained by this heterotic effect. Taken together, this makes the zebra finch a suitable model to study the link between sperm design and swimming speed and unravel the mechanism behind the relationship between inversion genotype, midpiece length and sperm velocity. Mendonca *et al.* [4] hypothesized that longer midpieces, known to contain more gyres, result from the fusion of a larger number of mitochondria with the potential to boost energy metabolism. Two recent studies using sperm morphology rather than the inversion genotype as their predictor did not fully support this hypothesis: mitochondrial volume did not increase linearly with midpiece length [4] and ejaculates of males with long midpieces contained less ATP molecules than those of males with shorter midpieces [15].

**Figure 1.**
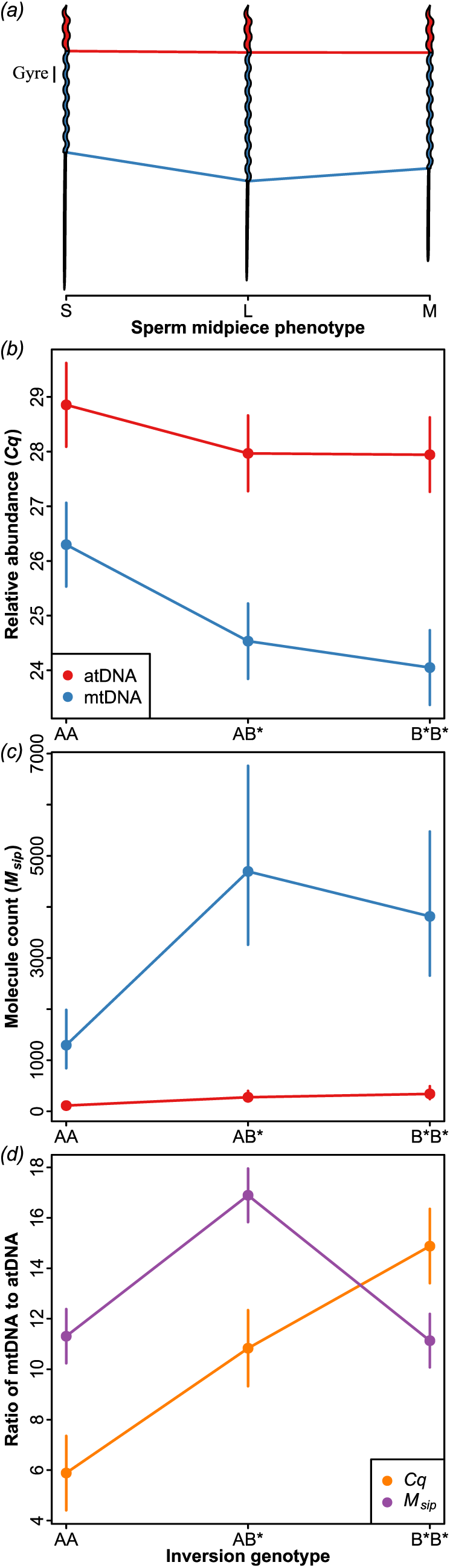
Effects of the chromosome *TguZ* inversion genotype on sperm phenotype (schematic) and on the amount (mean ± SE) of autosomal (nucleus, red) and mitochondrial (midpiece, blue) DNA in zebra finch ejaculates. AA and B*B* individuals are homozygous for the ancestral and derived inversion haplotypes, respectively. AB* individuals are heterozygous and have the longest midpieces. (*a*) Inversion genotype explained 41.9% of the variance in sperm midpiece length [2]. A single gyre is indicated. Effects were either measured as relative abundance *Cq*-values (*b*) or converted to molecule counts *M*_*sip*_ (*c*) and were transformed into a ratio of mitochondrial to autosomal DNA (*d*, orange for *Cq*-values and purple for *M*_*sip*_). Because a lower relative abundance *Cq*-value indicates more template DNA molecules, mirroring (*b*) on the x-axis should result in (*c*).

Here, we specifically test whether the inversion genotype has an effect on mitochondrial content in zebra finch ejaculates. We predict that the sperm of heterozygous males contain more copies of mitochondrial DNA than those of homozygous individuals.

## 2. Material & Methods

### Study subjects & sperm sampling

We extracted ejaculates by abdominal massage from 36 captive zebra finches (*Taeniopygia guttata castanotis*, average ± SD age at sampling = 6.3 ± 0.3 years) housed in a unisex group at the Max Planck Institute for Ornithology, Seewiesen, Germany. Birds were derived from study population “Seewiesen-GB” (see [16]), in which the connection between inversion genotype, sperm morphology and siring success had been previously established. Ejaculates were collected in 10 μl TNE buffer (10 mM Tris-HCl, 150 mM NaCl and 1 mM EDTA, pH 7.4), immediately frozen on dry ice and stored at −80°C until further usage.

### Inversion genotypes

All individuals had been previously genotyped for six SNPs that unambiguously tagged the three *TguZ* inversion haplotypes (named A, B and C) when using an unanimity decision rule (that is, all tag SNPs must specify the same type and missing data are not allowed; [12]). Further details on the genotyping and filtering procedure can be found elsewhere [12, 17]. In a previous study, Knief *et al.* [2] have shown that haplotype A represents the ancestral state and that haplotypes B and C have similar effects on sperm morphology and sperm velocity. Thus, as in Knief *et al.* [2], we here combine them into the derived haplotype B*. After removing one individual with low sperm DNA concentration, the final sample consisted of 9 individuals homozygous for the ancestral inversion haplotype (AA), 13 heterozygotes and 13 individuals homozygous for the derived inversion type (B*B*).

### Sperm DNA isolation

We used a custom protocol from Macherey-Nagel containing the GuEX buffer (50 mM Guanidine HCl, 10.5 mM Tris pH 8.0, 10.5 mM NaCl, 10.5 mM EDTA pH 8.0, 1 mM NaOH, pH 8.0–8.5) and Proteinase K together with the DNeasy Blood & Tissue Kit (Qiagen) for DNA isolation from whole ejaculates. The detailed protocol is provided in the **Supplement**.

### Quantitative real-time PCR

To estimate the ratio of autosomal to mitochondrial DNA in zebra finch sperm, we used quantitative real-time PCR (qPCR; see also [18-23]). All qPCR reactions were set up with the Luna Universal qPCR master mix (New England BioLabs) and 150 nM of each primer in a total volume of 20 μl. We used three autosomal primer pairs that had been previously assayed for their amplification efficiency in the zebra finch [24] and designed three mitochondrial primer pairs in the *ND2, ND4* and *ND5* genes (**Table S1**). We mixed DNA isolated from zebra finch liver and blood for normalization and tested the amplification efficiency of all primer pairs through a six- and seven-step log_10_ serial dilution of this standard (0.125–10 ng DNA for autosomal and 0.008–1 ng DNA for mitochondrial markers). All standard DNA samples were run in triplicates on the C1000 Touch™ Thermo Cycler with the optical module CFX96 Touch™ Real-Time PCR Detection System (Bio-Rad). An initial 5 min. denaturation step at 95°C was followed by 40 cycles of 15 sec. at 95°C and 30 sec. at 62°C and a final melt curve measurement from 65°C to 95°C. We obtained raw *Cq*-values through the Bio-Rad CFX Manager software (v3.1.1517.0823). All primer pairs had efficiency values between 97% and 110% (**Table S2**).

For the actual quantification of mitochondrial and autosomal DNA in sperm, we followed the same protocol as above, using 1 ng of sperm DNA as our template in every qPCR reaction. To estimate the run-specific amplification efficiency, we set up a three-step dilution series covering the sperm DNA quantities, starting from 5 ng and 1 ng standard DNA template for the autosomal and mitochondrial primer pairs, respectively. We attempted to run each sample in triplicate for one autosomal and one mitochondrial marker in every qPCR run, along with the dilution series of the standard DNA. In total, we amplified every sample with 7–8 combinations of the autosomal and mitochondrial primer pairs.

We excluded 7 qPCR reactions of two individuals that were outliers in terms of their *Cq*-values (more than 4 standard deviations from the mean *Cq*-value of that run). These were most likely technical artefacts because the same samples were amplified normally with the same primer pairs in different reactions. The decision to remove extreme outliers was taken blind to the outcome of the study. After this removal, we kept the data from 1494 PCR reactions from 24 qPCR runs. In this final data set, each of 35 individuals was genotyped within each run with each of the two primer pairs on average 3 times (range 1–6 times) and in total 7.3 times (range 1–12) with every primer pair (*N* = 6 primer pairs).

### Statistical analyses

For each qPCR run *i* and each primer pair *p*, we estimated the efficiency (*E*_*ip*_) by fitting a linear regression model of *Cq*-values on the log_10_ DNA-concentrations of the standard amplification curve. We took the slope *β*_*ip*_ of this regression to calculate the efficiency as

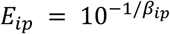

Ideally, exponential amplification leads to a doubling of the number of DNA molecules in every cycle of the PCR, which translates into an efficiency value of 2. This value can be converted into a percentage (*EP*_*ip*_) by calculating

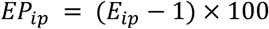

We used two different approaches for correcting the raw *Cq*-values (*Cq*_*sip*_) of each sample (*s*) for the amplification efficiency of the primer pair (*p*) used in each qPCR run (*i*). First, we calculated the relative abundance (*R*_*sip*_) as described in Steibel *et al.* [25]:

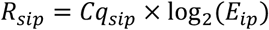

Alternatively, we transformed the raw *Cq*-values into molecule counts (*M*_*sip*_) as described in Matz *et al.* [26]:

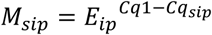

where *Cq1* is the *Cq*-value when using a single DNA molecule as PCR template. It was either set to 37 (see [26]) or estimated from the standard curve of each primer pair of every qPCR run. Because there was no qualitative difference, we present results in which we used empirical *Cq1*-values (*Cq1*_*ip*_).

We then fitted mixed-effects linear models with efficiency corrected *Cq*-values (*R*_*sip*_ or *M*_*sip*_) as the dependent variables. To test our hypothesis that sperm with longer midpieces (genotype AB*) contain more copies of the mitochondrial DNA, we fitted the interaction between the two independent variables “chromosome type” (that is autosome vs mitochondrion, 1 degree of freedom) and “inversion genotype” (2 degrees of freedom, i.e. genotype AA with short, genotype B*B* with intermediate and genotype AB* with long sperm midpieces). In both models, we corrected for individual ID (35 levels), primer pair (6 levels) and qPCR run (24 levels) by fitting them as random effects. Using *R*_*sip*_ as the dependent variable, we fitted a model with a Gaussian error structure and all of the above fixed and random effects. We further analysed *M*_*sip*_ by fitting a generalized linear mixed-effects model with a Poisson error structure. To account for overdispersion [27], we added an observation-level random effect (OLRE; [28]). Primer pair did not explain any variation in *M*_*sip*_ and was dropped from the final model.

We also estimated the difference between autosomal and mitochondrial DNA content with the familiar Δ*Cq*-method [29] which assumes efficiency values of 2 (= 100%) for every marker and run. To this end, we first calculated the mean *Cq*-value for every sample and marker within each run, then subtracted the mean mitochondrial *Cq*-value from the mean autosomal *Cq*-value of every sample within each run and fitted a linear regression model with “inversion genotype” as the sole predictor. Effects were in the same direction as those obtained with the mixed-effects model using *R*_*sip*_ as the dependent variable and fitting “inversion genotype” was also highly significant (*P* = 0.006). Because mixed-effects models are more flexible and allow controlling for additional random variation [25], we present results of the mixed-effects models only.

All analyses were conducted using R (v3.4.3; [30]). Linear mixed-effects and generalized linear mixed-effects models were fitted with the lmer() and glmer() function of the lme4 (v1.1-21; [31]), respectively. Contrasts and *P*-values were estimated with the lmerTest package (v3.1-0; [32]). Standard errors of ratios of fixed effects were derived through bootstrapping using the bootMer() function with 1000 resampled data sets. Some models failed to converge during the bootstrapping procedure, but the bootstrapped fixed effect estimates and their standard errors were equivalent to those obtained through lmer() and glmer() up to the second decimal. Raw data and analyses scripts are available as **Supplementary data**.

## 3. Results

As expected, zebra finch ejaculates contained significantly more mitochondrial (mtDNA) than autosomal DNA (atDNA) copies (*P* = 3 × 10^−4^ using relative abundance and *P* < 2 × 10^−16^ using molecule counts). Excluding the interaction between chromosome type and inversion genotype from the statistical model and using either the relative abundance or the molecule count data, ejaculates contained 10.5 ± 1.5 (expressed as a ratio ± SE) and 12.9 ± 1.0 times more copies of mitochondrial than autosomal DNA.

The amount of mtDNA relative to the amount of atDNA differed between males, depending on their *TguZ* inversion genotype. Males producing sperm with short midpieces (genotype AA) had less mitochondrial DNA in their ejaculates than those with intermediate or long midpieces (genotypes B*B* and AB*, respectively; interaction *P* = 3 × 10^−15^ using relative abundance and *P* = 1 × 10^−7^ using molecule counts, **Figure 1b,c**). In the relative abundance data, the amount of mtDNA seemed to increase linearly with each copy of the derived inversion type B* (ratio ± SE of mtDNA to atDNA, AA: 5.9 ± 1.5, AB*: 10.8 ± 1.5, B*B*: 14.9 ± 1.5; **Figure 1d, Table 1**). In contrast, mtDNA molecule counts suggested that sperm with the longest midpieces (genotype AB*) had the highest mtDNA to atDNA ratio (ratio ± SE of mtDNA to atDNA, AA: 11.3 ± 1.1, AB*: 16.9 ± 1.1, B*B*: 11.1 ± 1.1; **Figure 1d, Table 1**).

**Table 1.**
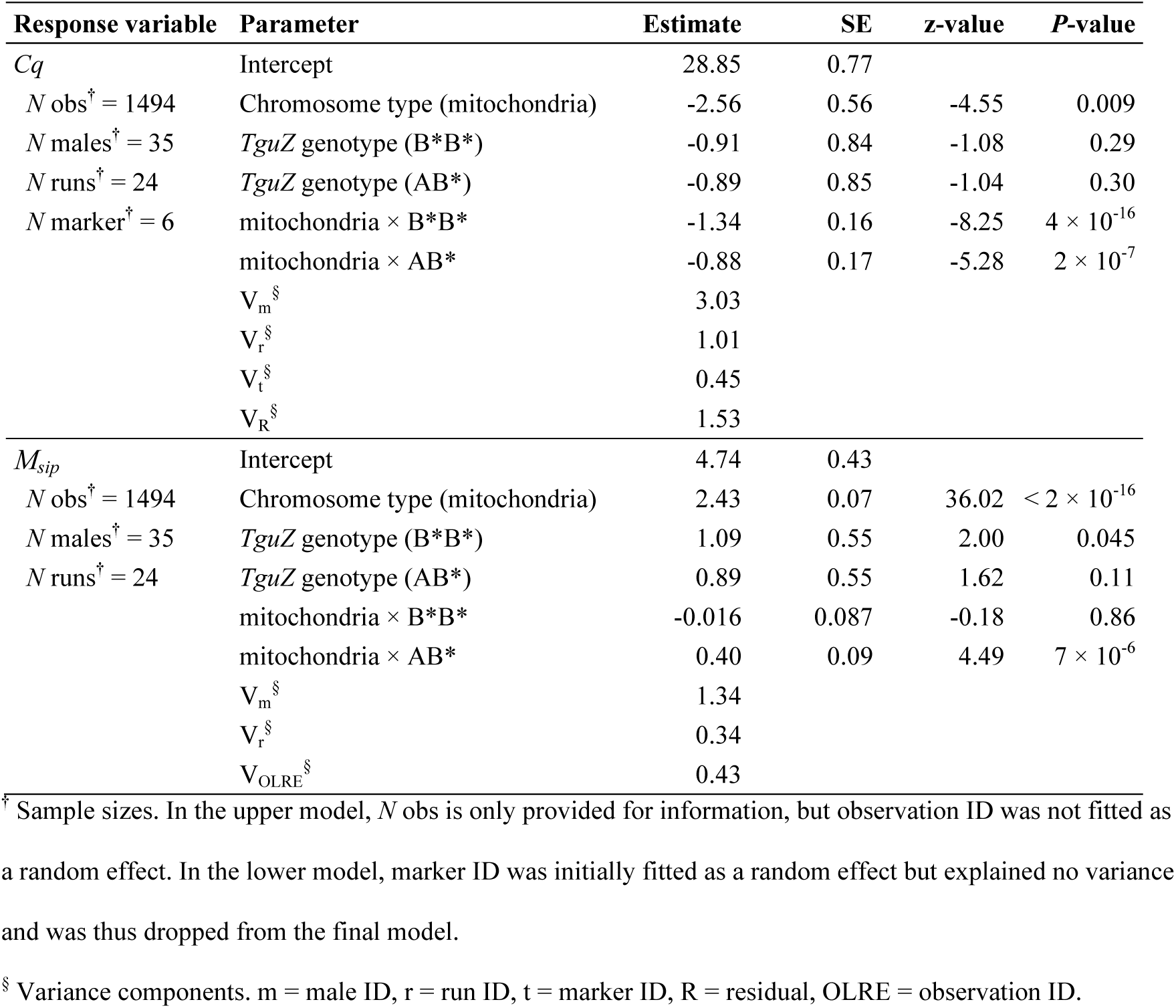
Estimates from the full mixed-effects models using either relative abundance (*Cq*) or molecule counts (*M*_*sip*_) as the response variable. Because the molecule counts model had a Poisson error structure with a log-link function, all parameter estimates are on the log-scale. There was no residual variance component estimated and the observation-level random effect (V_OLRE_) represents the residual.

## Discussion

Here, we have shown that zebra finch ejaculates contain about 10–13 times more copies of mitochondrial than autosomal DNA. The magnitude of this difference was significantly affected by the inversion genotype of the male, such that heterozygous males have more mtDNA copies than those homozygous for the ancestral and — depending on the analyses — also the derived inversion haplotype. Since heterozygous males have the longest sperm midpieces [2, 14] this difference is consistent with the hypothesized relationship between sperm morphology and energy metabolism. Although we did not directly measure sperm midpiece length in the sampled ejaculates, the inversion genotype is a reliable predictor of sperm midpiece length in this zebra finch population. The inversion genotype explains some 41.9% of the variation in midpiece length and males heterozygous for the inversion have on average 26.59% and 11.16% longer midpieces than males homozygous for the ancestral and derived genotypes, respectively [2]. It should be noted that ejaculates usually also contain small amounts of non-sperm cells [33], such that the inversion genotype could in principle also affect the cellular composition of ejaculates. It would, however, require large mtDNA copy numbers in these few cells to significantly bias our estimated ratio of mitochondrial to autosomal DNA upwards.

In humans, a normal sperm cell contains about 15 mitochondria [34] and across mammalian tissues, there seems to be a consistent number of 1–3 mtDNA copies per mitochondrion ([35, 36]; but [37] estimate 1–10 copies). Using qPCR, the average mtDNA copy number for a single human sperm cell is 11.13 (range = 0.2–344.9, *N* = 795 males), which translates roughly into one mtDNA copy per mitochondrion in every sperm cell [18-23]. Assuming the same to be true in passerines, then about 10–13 mitochondria could contribute to the mitochondrial syncytium of a single sperm midpiece in zebra finches, ranking lowest in comparison to other avian species (15–350 mitochondria per sperm cell; [6]). In oscine songbirds, the mitochondrial syncytium winds along the midpiece (gyres) and a single gyre in zebra finch is 3.783 μm long [4]. Taking the average midpiece lengths for the three inversion genotypes [2], short midpieces have 6.55 gyres (genotype AA), intermediate 7.46 (genotype B*B*) and long midpieces 8.29 (genotype AB*, **Figure 1a**). Using the average of the relative abundance and molecule count estimates, these morphological measurements correlate well with the predicted number of 8.6 (genotype AA), 13.0 (genotype B*B*) and 13.9 (genotype AB*) mitochondria. This would mean that every gyre is stably composed of roughly 1.6 (AA: 1.3, B*B*: 1.7, AB*: 1.7) mitochondria regardless of genotype. Taking estimates from Mendonca *et al.* [4], individuals producing sperm with the longest midpieces (genotype AB*) have about 14% larger sperm midpiece volumes than individuals of genotype AA (V = 2.4 and 2.1 μm^3^ for AB* and AA, respectively). Yet, the number of mitochondria increases by 61 % (*N* = 13.9 and 8.6). This is similar to the 44 % increase in midpiece length (mean length = 31.4 and 21.8 μm [4]) and the number of gyres (8.30 and 5.76 [4]) suggesting that the number of mitochondria in the syncytium is primarily determined by midpiece length rather than volume.

Yet, why should a larger number of mtDNA molecules increase the rate of ATP production, considering that only 13 proteins of the respiratory chain are encoded in the mitochondrial DNA and the remaining roughly 100 proteins in the nuclear DNA [38]? In non-sperm tissue it has been shown that mtDNA copy number is correlated with respiratory activity [39] and sexually dimorphic gene expression [40], and transcript levels of the protein-coding mitochondrial genes in human sperm covary with mtDNA copy number [41]. Furthermore, mitochondrial ribosomes have been shown to be translationally active in sperm cells; inhibiting them reduces sperm motility [42]. Thus, a larger number of mtDNA copies could directly affect the number of mitochondrial ribosomes, either because mtDNA copy number serves a proxy for the amount of mitochondrial ribosomes or because more mtDNA copies would allow higher transcription rates of their ribosomal RNAs [43].

In contrast to our results, human and other mammalian males with pathologic sperm diseases have more mtDNA copies in their sperm cells than healthy males and sperm with more mtDNA copies swim slower ([18, 20, 44, 45] and references therein, but see [46, 47] for the opposite finding). This is often explained by larger quantities of reactive oxygen species (ROS) produced in sperm with more mitochondria, which could damage the sperm cells [48]. However, most studies were clinical and not experimental, such that the causal relationship has not been fully resolved. Males might actually compensate for high amounts of abnormal sperm by harbouring more mtDNA copies in each sperm cell, which has been found to restore fertility [49]. Furthermore, human sperm cells seem to derive energy also through anaerobic glycolysis in the tail, such that OXPHOS might be unnecessary for motility [5]. The glycolytic enzymes are located along a fibrous sheath, a structure that is seemingly absent in passerine sperm cells [4]. Thus, findings extracted for humans may qualitatively differ from avian sperm physiology.

Summarizing, this study establishes an intriguing link between naturally segregating inversion genotypes, sperm morphology and physiology and motivates future research of energy metabolism in male gametes of non-mammalian models.

## Ethics

Housing, breeding, blood and sperm sampling of the captive zebra finches are covered by the housing and breeding permit granted to W.F. (# 311.4-si, Landratsamt Starnberg, Germany).

## Data accessibility

Raw qRT-PCR data and analyses scripts are available as **Supplementary data**.

## Author’s contribution

U.K. and J.B.W.W. conceived of the study. U.K. performed research and analysed the data. W.F. and B.K. provided samples and intellectual input. U.K. wrote the paper with input from all authors.

## Competing interests

The authors declare no competing interests.

## Funding

Funding was provided by the LMU Munich (to J.B.W.W.) and the Max Planck Society (to B.K.).

## Acknowledgements

We thank K. Martin for help with breeding and sperm sampling and S. Bauer, E. Bodendorfer, A. Grötsch, A. Kortner, K. Martin, P. Neubauer, F. Weigel and B. Wörle for animal care. F. Martínez-Pastor provided the TNE buffer recipe. We are further grateful to G. Kumpfmüller for laboratory work and thank J. Peñalba for the sperm illustrations.

## References

1. Bennison C, Hemmings N, Slate J, Birkhead T. Long sperm fertilize more eggs in a bird. P R Soc B. 2015;282(1799):e20141897.

2. Knief U, Forstmeier W, Pei YF, Ihle M, Wang DP, Martin K, et al. A sex-chromosome inversion causes strong overdominance for sperm traits that affect siring success. Nat Ecol Evol. 2017;1(8):1177–84.

3. Parker GA. Sperm competition and the evolution of ejaculates: towards a theory base. In: Birkhead TR, Møller AP, editors. Sperm competition and sexual selection. London, UK: Academic Press; 1998. p. 3–54.

4. Mendonca T, Birkhead TR, Cadby AJ, Forstmeier W, Hemmings N. A trade-off between thickness and length in the zebra finch sperm mid-piece. P R Soc B. 2018;285(1883):e20180865.

5. Cummins J. Sperm motility and energetics. In: Birkhead TR, Hosken DJ, Pitnick S, editors. Sperm biology: an evolutionary perspective. London, UK: Elsevier; 2009. p. 185–206.

6. Jamieson BGM. Avian spermatozoa: structure and phylogeny. In: Jamieson BGM, editor. Reproductive biology and phylogeny of birds. 6A. Enfield, NH: Science Publisher; 2007. p. 349–511.

7. Froman DP, Feltmann AJ. Sperm mobility: a quantitative trait of the domestic fowl (*Gallus domesticus*). Biol Reprod. 1998;58(2):379–84.

8. Fawcett DW, Anderson WA, Phillips DM. Morphogenetic factors influencing the shape of the sperm head. Dev Biol. 1971;26(2):220–51.

9. Vernon GG, Woolley DM. Three-dimensional motion of avian spermatozoa. Cell Motil Cytoskeleton. 1999;42(2):149–61.

10. Góes RM, Dolder H. Cytological steps during spermiogenesis in the house sparrow (*Passer domesticus*, Linnaeus). Tissue & Cell. 2002;34(4):273–82.

11. Rowe M, Laskemoen T, Johnsen A, Lifjeld JT. Evolution of sperm structure and energetics in passerine birds. P R Soc B. 2013;280(1753):e20122616.

12. Knief U, Hemmrich-Stanisak G, Wittig M, Franke A, Griffith SC, Kempenaers B, et al. Fitness consequences of polymorphic inversions in the zebra finch genome. Genome Biology. 2016;17:e199.

13. Itoh Y, Kampf K, Balakrishnan CN, Arnold AP. Karyotypic polymorphism of the zebra finch Z chromosome. Chromosoma. 2011;120(3):255–64.

14. Kim KW, Bennison C, Hemmings N, Brookes L, Hurley LL, Griffith SC, et al. A sex-linked supergene controls sperm morphology and swimming speed in a songbird. Nat Ecol Evol. 2017;1(8):1168–76.

15. Bennison C, Hemmings N, Brookes L, Slate J, Birkhead T. Sperm morphology, adenosine triphosphate (ATP) concentration and swimming velocity: unexpected relationships in a passerine bird. P R Soc B. 2016;283(1837):e20161558.

16. Forstmeier W, Segelbacher G, Mueller JC, Kempenaers B. Genetic variation and differentiation in captive and wild zebra finches (*Taeniopygia guttata*). Mol Ecol. 2007;16(19):4039–50.

17. Knief U, Schielzeth H, Backström N, Hemmrich-Stanisak G, Wittig M, Franke A, et al. Association mapping of morphological traits in wild and captive zebra finches: reliable within, but not between populations. Mol Ecol. 2017;26(5):1285–305.

18. Amaral A, Ramalho-Santos J, St John JC. The expression of polymerase gamma and mitochondrial transcription factor A and the regulation of mitochondrial DNA content in mature human sperm. Hum Reprod. 2007;22(6):1585–96.

19. May-Panloup P, Chretien MF, Savagner F, Vasseur C, Jean M, Malthièry Y, et al. Increased sperm mitochondrial DNA content in male infertility. Hum Reprod. 2003;18(3):550–6.

20. Song GJ, Lewis V. Mitochondrial DNA integrity and copy number in sperm from infertile men. Fertil Steril. 2008;90(6):2238–44.

21. Zhang GW, Wang Z, Ling X, Zou P, Yang H, Chen Q, et al. Mitochondrial biomarkers reflect semen quality: results from the MARCHS study in Chongqing, China. Plos One. 2016;11(12):e0168823.

22. Tian MP, Bao HQ, Martin FL, Zhang J, Liu LP, Huang QY, et al. Association of DNA methylation and mitochondrial DNA copy number with human semen quality. Biol Reprod. 2014;91(4):e101.

23. Wu HT, Huffman AM, Whitcomb BW, Josyula S, Labrie S, Tougias E, et al. Sperm mitochondrial DNA measures and semen parameters among men undergoing fertility treatment. Reprod Biomed Online. 2019;38(1):66–75.

24. Wolf JBW, Bryk J. General lack of global dosage compensation in ZZ/ZW systems? Broadening the perspective with RNA-seq. Bmc Genomics. 2011;12:e91.

25. Steibel JP, Poletto R, Coussens PM, Rosa GJ. A powerful and flexible linear mixed model framework for the analysis of relative quantification RT-PCR data. Genomics. 2009;94(2):146–52.

26. Matz MV, Wright RM, Scott JG. No control genes required: Bayesian analysis of qRT-PCR data. Plos One. 2013;8(8):e71448.

27. Knief U, Forstmeier W. Violating the normality assumption may be the lesser of two evils. bioRxiv. 2018.

28. Harrison XA. Using observation-level random effects to model overdispersion in count data in ecology and evolution. Peerj. 2014;2:e616.

29. Livak KJ, Schmittgen TD. Analysis of relative gene expression data using real-time quantitative PCR and the 2-^ΔΔCT^ method. Methods. 2001;25(4):402–8.

30. R Core Team. R: a language and environment for statistical computing. 3.4.3 ed. Vienna, Austria: R Foundation for Statistical Computing; 2017.

31. Bates D, Mächler M, Bolker BM, Walker SC. Fitting linear mixed-effects models using lme4. J Stat Softw. 2015;67(1):1–48.

32. Kuznetsova A, Brockhoff PB, Christensen RHB. lmerTest package: tests in linear mixed effects models. J Stat Softw. 2017;82(13):1–26.

33. Fedder J. Nonsperm cells in human semen: with special reference to seminal leukocytes and their possible influence on fertility. Arch Andrology. 1996;36(1):41–65.

34. Bedford JM, Hoskins DD. The mammalian spermatozoon: morphology, biochemistry and physiology. In: Lamming GE, editor. Marshall’s physiology of reproduction: Reproduction in the male. 2. 4 ed. London: Churchill Livingstone; 1990. p. 379–568.

35. Wiesner RJ, Rüegg JC, Morano I. Counting target molecules by exponential polymerase chain reaction: copy number of mitochondrial DNA in rat tissues. Biochem Bioph Res Co. 1992;183(2):553–9.

36. Robin ED, Wong R. Mitochondrial DNA molecules and virtual number of mitochondria per cell in mammalian cells. J Cell Physiol. 1988;136(3):507–13.

37. Satoh M, Kuroiwa T. Organization of multiple nucleoids and DNA molecules in mitochondria of a human cell. Exp Cell Res. 1991;196(1):137–40.

38. Rantanen A, Larsson NG. Regulation of mitochondrial DNA copy number during spermatogenesis. Hum Reprod. 2000;15 Suppl 2:86–91.

39. D’Erchia AM, Atlante A, Gadaleta G, Pavesi G, Chiara M, De Virgilio C, et al. Tissue-specific mtDNA abundance from exome data and its correlation with mitochondrial transcription, mass and respiratory activity. Mitochondrion. 2015;20:13–21.

40. Camus MF, Wolf JBW, Morrow EH, Dowling DK. Single nucleotides in the mtDNA sequence modify mitochondrial molecular function and are associated with sex-specific effects on fertility and aging. Curr Biol. 2015;25(20):2717–22.

41. Pietropaolo V, Passariello C, Bellizzi A, Virga A, Anzivino E, Rodio DM, et al. Analysis of sperm motility related to transcriptional alterations of mitocondrial genes in males affected by infertility. Eur J Inflamm. 2012;10(3):455–62.

42. Gur Y, Breitbart H. Mammalian sperm translate nuclear-encoded proteins by mitochondrial-type ribosomes. Gene Dev. 2006;20(4):411–6.

43. de Paula WB, Agip AN, Missirlis F, Ashworth R, Vizcay-Barrena G, Lucas CH, et al. Female and male gamete mitochondria are distinct and complementary in transcription, structure, and genome function. Genome Biol Evol. 2013;5(10):1969–77.

44. Wu HT, Whitcomb BW, Huffman A, Brandon N, Labrie S, Tougias E, et al. Associations of sperm mitochondrial DNA copy number and deletion rate with fertilization and embryo development in a clinical setting. Hum Reprod. 2019;34(1):163–70.

45. Moraes CR, Meyers S. The sperm mitochondrion: organelle of many functions. Anim Reprod Sci. 2018;194:71–80.

46. Mundy AJ, Ryder TA, Edmonds DK. Asthenozoospermia and the human sperm mid-piece. Hum Reprod. 1995;10(1):116–9.

47. Kao SH, Chao HT, Liu HW, Liao TL, Wei YH. Sperm mitochondrial DNA depletion in men with asthenospermia. Fertil Steril. 2004;82(1):66–73.

48. Tremellen K. Oxidative stress and male infertility — a clinical perspective. Hum Reprod Update. 2008;14(3):243–58.

49. Jiang M, Kauppila TES, Motori E, Li XP, Atanassov I, Folz-Donahue K, et al. Increased total mtDNA copy number cures male infertility despite unaltered mtDNA mutation load. Cell Metab. 2017;26(2):429–36.

