## Supplementary Material for "A sex chromosome inversion is associated with copy number variation of mitochondrial DNA in zebra finch sperm"

#### Index

**Supplementary Methods** 2
**Supplementary Tables** 3

### Supplementary Methods

**Sperm DNA extraction**

We used a custom protocol from Macherey-Nagel containing the GuEX buffer (50 mM Guanidine HCl, 10.5 mM Tris pH 8.0, 10.5 mM NaCl, 10.5 mM EDTA pH 8.0, 1 mM NaOH, pH 8.0–8.5) and Proteinase K together with the DNeasy Blood & Tissue Kit (Qiagen) for DNA isolation from whole ejaculates. The detailed protocol was:

1. Mix the whole ejaculate with 475 µl GuEX + 25 µl Proteinase K (20 mg/µl, Macherey-Nagel).
2. Incubate for 15 min. at 37°C.
3. Then centrifuge at 12,000 g for 5 minutes at room temperature.
4. Resuspend pellet in 350 µl GuEX and centrifuge at 12,000 g for 5 minutes at room temperature.
5. Repeat step (3) 2–3 times.
6. Resuspend pellet in 180 µl buffer ATL (Qiagen), vortex and add 20 µl Proteinase K (20 mg/µl, Macherey-Nagel).
7. Incubate overnight at 60°C.
8. Add 4 µl RNase A (100 mg/µl).
9. Incubate for 10 min. at 37°C.
10. Follow the protocol: Purification of total DNA from animal blood or cells (spin-column protocol) of the Qiagen DNeasy Blood & Tissue Kit (steps 2–8) but elute with 25 µl and 20 µl EB (10 mM Tris, pH 8.5).

DNA quantity was assessed using the Qubit (Invitrogen).

### Supplementary Tables

**Table S1 |** Descriptive summary of the six primer pairs used. The first three primer pairs amplify DNA from the autosomes (*Tgu1*, *Tgu5* and *Tgu27*) and were published previously in Wolf and Bryk [1]. The last three amplify mitochondrial DNA (*TguM*).

| Chromosome | Gene ID | Gene name | Gene region | Internal primer code | Forward primer | Reverse primer | Amplicon length (bp) |
|---|---|---|---|---|---|---|---|
| *Tgu1* | ENSTGUG00000013338 | *GAPDH* | exon 7 | A12 | CACACAGAAGACAGTGGATGG | ACTTTTCCCACAGCCTTAGC | 111 |
| *Tgu5* | ENSTGUG00000012606 | *CHGA* | exon 6 | A8 | AGGATGTGAGAAGGTCATGG | CTTCTTTTCTTCAGGCAtGG | 89 |
| *Tgu27* | ENSTGUG00000002022 | *NSF* | exon 12 | A1 | TCCAGATCCTGCACATCC | GCTCGAACCAAACCTTCC | 130 |
| *TguM* | ENSTGUG00000018759 | *ND4* | exon1 | M2 | CTAGTCGTAGCCGCAACAAT | GCTGTGGGTTCGTTCATAGT | 132 |
| *TguM* | ENSTGUG00000018763 | *ND5* | exon1 | M3 | ACCCTTCTAGCCACATCCTT | TTACTGCGGGGTTGTTTTCA | 118 |
| *TguM* | ENSTGUG00000018741 | *ND2* | exon1 | M5 | AGTCCGAAAAGTCCTAGCCT | TGAGTTTGGGGTTGTACGTG | 80 |

**Table S2 |** Amplification efficiency of the six primer pairs using the standard DNA as template.

| Internal primer code | Regression slope β | Multiple $R^2$ | Efficiency (%) |
|---|---:|---:|---:|
| A12 | -3.104 | 0.986 | 110.0 |
| A8 | -3.195 | 0.988 | 105.6 |
| A1 | -3.270 | 0.993 | 102.2 |
| M2 | -3.379 | 0.995 | 97.7 |
| M3 | -3.318 | 0.991 | 100.2 |
| M5 | -3.287 | 0.997 | 101.5 |

#### References

1. Wolf JBW, Bryk J. General lack of global dosage compensation in ZZ/ZW systems? Broadening the perspective with RNA-seq. Bmc Genomics. 2011;12:e91.
